# Equine herpesvirus myeloencephalopathy-related mutations in EHV-1 DNA polymerase allow EHV-1 to grow at elevated temperatures

**DOI:** 10.1101/2025.08.22.671711

**Authors:** Noriko Fukushi, Rikio Kirisawa, Fuka Nishimura, Koji Tsujimura, Hideto Fukushi

## Abstract

Equine herpesvirus myeloencephalopathy (EHM) caused by equine herpesvirus type 1 (EHV-1) is a major threat to the equine industry because of its devastatating impact on animal welfare and performance. A specific single nucleotide polymorphism (SNP) (G2254/D752) in the EHV-1 ORF30 gene encoding DNA polymerase was previously shown to be a marker of the strain of EHV-1 that causes EHM. However, the effect of this SNP has never been resolved. Clinical findings indicated that fever and higher viremia levels are associated with the onset of EHM. No studies have examined EHV-1 growth at elevated temperatures so far. We found that EHV-1 with the D752 SNP replicated at elevated temperatures, and EHV-1 without it did not. We also found that EHV-1s isolated from horses with EHM could grow at elevated temperatures and most non-EHM isolates were suppressed by elevated temperatures. EHV-1s that can replicate at elevated temperatures have this SNP or one other SNP in ORF30. This appears to be the first report to show an association between EHM and the ability of EHV-1 to grow at elevated temperatures. (177 words)

**Importance:** Equine herpesvirus myeloencephalopathy (EHM) caused by equine herpesvirus type 1 (EHV-1) is a major threat to the equine industry because of its impact on animal welfare and performance. A specific single nucleotide polymorphism (SNP) (G2254/D752) in EHV-1 ORF30 gene encoding DNA polymerase (UL30) was previously shown to be a marker of EHM. However, the effect of this SNP has never been resolved. Here we show that this and one other SNP in UL30 are associated with replication capacity at elevated temperatures. (81 words)

## Introduction

Equine herpesvirus type 1 (EHV-1: family *Orthoherpesviridae*, subfamily *Alphaherpesvirinae*, *Varicellovirus equidalpha1*) causes respiratory infection, abortion and neurological disease in horses worldwide (1–3). It can lead to a neurological disease called equine herpesvirus myeloencephalopathy (EHM), which can damage the spinal cord and brain. The incidence of EHM has been increasing in Europe and the United States in recent years (4). Disability may remain after recovery from EHM (5). Following an outbreak in Valencia, Spain in 2021 at an international event, many exposed horses were transported back to their respective countries where they developed EHM (6, 7). Fever is one of the clinical signs of EHM (8–11). The onset of EHM occurs 1 to 3 days after resolution of a fever (7). Horses with EHM have longer periods of viremia and higher levels of virus in their blood (12, 13).

A single nucleotide polymorphism (SNP) (A2254G),which causes an N752D substitution in a DNA polymerase (UL30), has been implicated in EHM (14). Another SNP (C2254) causing an N752H substitution in UL30 has been observed in EHM isolates in France (15) and the United States (16). The neuropathogenic genotype G2254/D752 has been associated with various virus characteristics, such as fever and higher viremia load (17–19), although it was not reported to have any significant effects on clinical signs or disease outcome (20). Furthermore, some EHV-1 strains causing EHM have N at position 752 in UL30 (14). The role of the amino acid at position 752 in the neuropathogenicity of EHV-1 has been a subject of debate (21, 22). Goodman et al. reported that the SNP (D752) determined neuropathogenicity of EHV-1(18). On the other hand, residue 752 in UL30 was reported to be non-essential for virus growth in vitro (22). Further studies are needed understand the role of the G2254/D752 and C2254/H752 genotypes in EHM outbreaks.

Here, to clarify the role of D752, we examined its effect on the replication capacity of EHM-derived EHV-1 at elevated temperatures. We hypothesized that EHV-1 proliferates well in the respiratory tract (33°C in the nasal cavity and 38°C in the lungs) of horse, but that its proliferation is inhibited at higher temperatures (39°C or higher), i.e. at typical fever temperatures in infected horses. However, no studies examined EHV-1 growth at elevated temperatures so far. We hypothesized that some strains of EHV-1 could proliferate at elevated temperatures, leading to high viral loads, resulting in viral proliferation in vascular endothelial cells, which in turn could lead to EHM. In this study, we aimed to clarify the relationship between replication capacity of EHV-1 at elevated temperatures and the SNPs in ORF30 (the ORF that encodes UL30).

## Results

### Effect of elevated temperature on viral replication

We compared the effects of temperatures between 37.0°C and 40.5°C on the replications of three viruses (23): a parent virus (EHM-derived Ab4p UL30 D752), a mutant virus (Ab4p UL30 N752), and a repaired virus (Ab4p Repair UL30 D752) (23) with an N752D substitution. All three viruses formed plaques at 38.5°C but not at 40.5°C. The mutant virus (N752) did not form plaques at 39.0°C or higher. The other two viruses formed plaques up to 40.0°C. Plaque sizes of the parent and repair viruses at 38.5 to 39.5°C were the same as those at 37.0°C and smaller at 40.0°C (Fig. 1). Plaque sizes of the mutant virus at 38.5°C were smaller than those at 37.0°C. The numbers of plaques formed by the parent and repair viruses at 38.5°C and higher temperatures were almost the same as the number at 37.0°C, but the numbers of plaques formed by the mutant virus at 39.0°C and higher temperatures were about a million-fold less than the number at 37.0°C (Table 1). These results indicate that Ab4p viruses possessing UL30 D752 formed plaques at elevated temperatures but Ab4p virus possessing UL30 N752 did not.

**Fig. 1.**
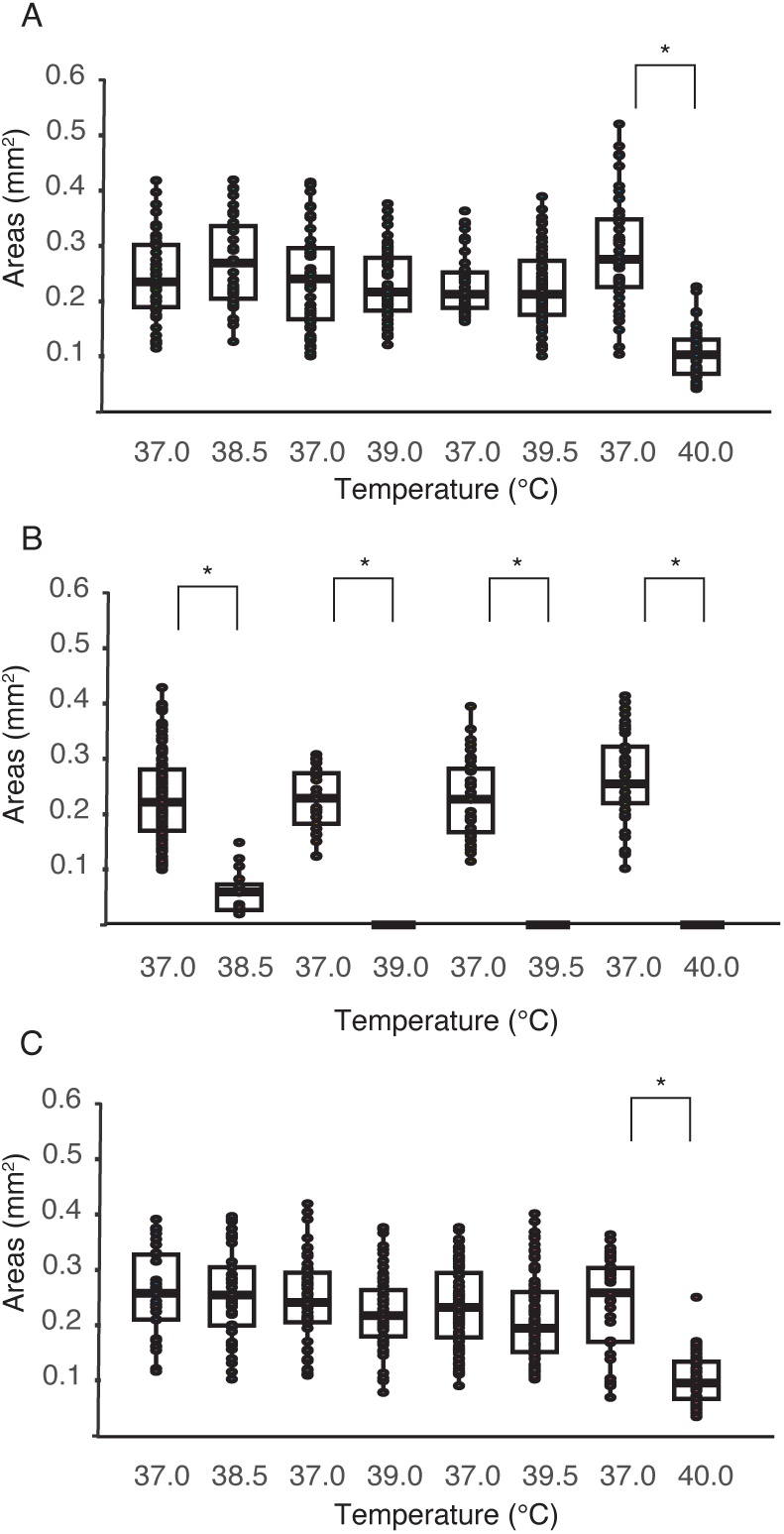
Plaque formation of Ab4p UL30 D752, UL30 N752, repair UL30 D752 viruses at elevated temperatures. Plaque assay was performed on FHK Tcl3.1 cells infected at 37°C or elevated temperature as indicated (38.5°C, 39.0°C, 39.5°C and 40.0°C) with Ab4p UL30 D752 (Parent) (A), Ab4p UL30 N752 (Mutant) (B) and Ab4p UL30 D752 (Repair) viruses (C). The graph shows areas of plaques formed by each virus and at each temperature. Boxplot is drawn with using ggplot package in RStudio version 2024.12.1+563. Dots show actual data. Data were compared using Wilcoxon rank sum test. P value is indicated as: *P < 0.05.

**Table 1.** Plating efficiency of the parent, mutant and repair viruses at elevated temperatures.

| Viruses | 38.5°C | 39.0°C | 39.5°C | 40.0°C |
| --- | --- | --- | --- | --- |
| Parent | 1.03 | 1.0 | 1.08 | 1.02 |
| Mutant | 0.63 | $\leq 10^{-6}$* | $\leq 10^{-6}$* | $\leq 10^{-6}$* |
| Repair | 1.0 | 1.21 | 1.0 | 1.07 |

The numbers indicate the plaque numbers at elevated temperature/the plaque numbers at 37°C
\*The number of plaques at elevated temperatures were under the detection limit.

The parental and repair viruses grew at 39.5°C, although their titers at 24 hours were 1/10 those at 37.0°C (Fig. 2). On the other hand, the mutant virus did not proliferate at 39.5°C. Real-time PCR targeting the ORF30 gene in infected cells showed that the copy numbers of the parent and the repair viruses at 39.5°C were the same as those at 37.0°C (Fig. 3 A, C). On the other hand, the copy number of the mutant at 39.5°C was at least 100-fold lower than at 37.0°C (Fig. 3 B). Replication of the mutant virus was reduced or halted at 39.5°C.

**Fig. 2.**
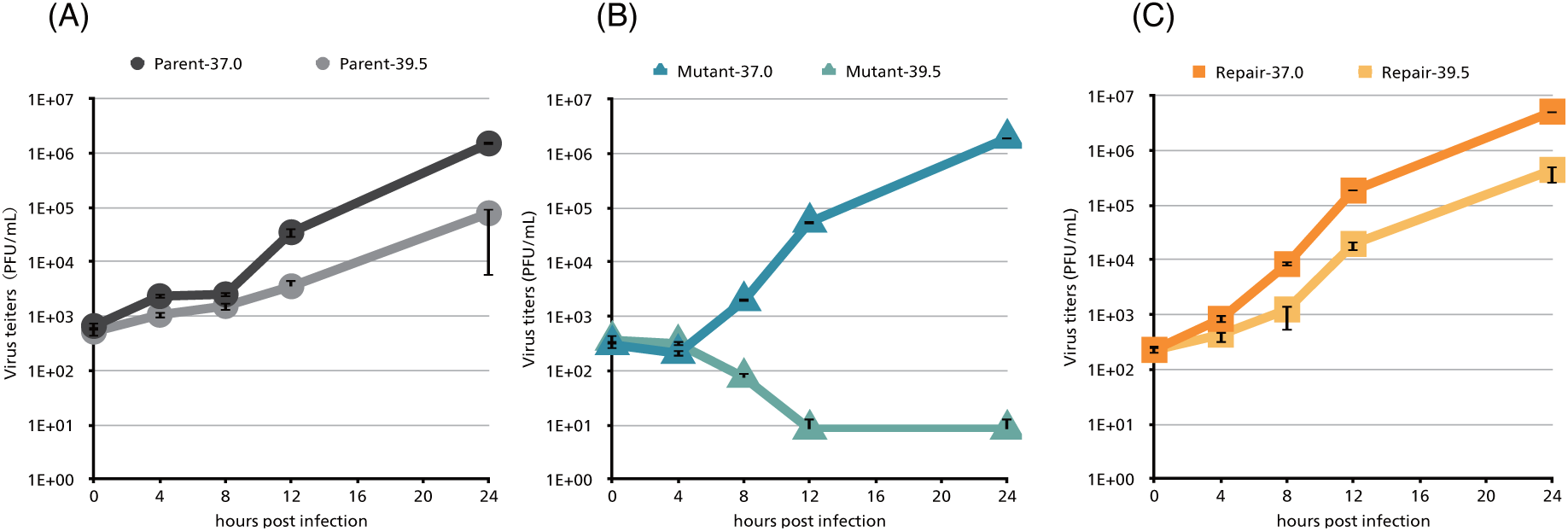
Growth kinetics of Ab4p viruses at 37.0℃ and 39.5℃. FHK Tcl3.1 cells were inoculated with virus and incubated at temperatures indicated in the figure. Culture supernatant was collected at 0, 4, 8, 12 and 24 hours post infection. Virus titers were assayed by plaque formation. (A) parent Ab4p virus, (B) mutant Ab4p ORF30 N752 virus, (C) repair Ab4p ORF30 D752 virus. Ab4p parent and repair viruses grew at 39.5°C. Mutant did not grow at 39.5°C.

**Fig. 3.**
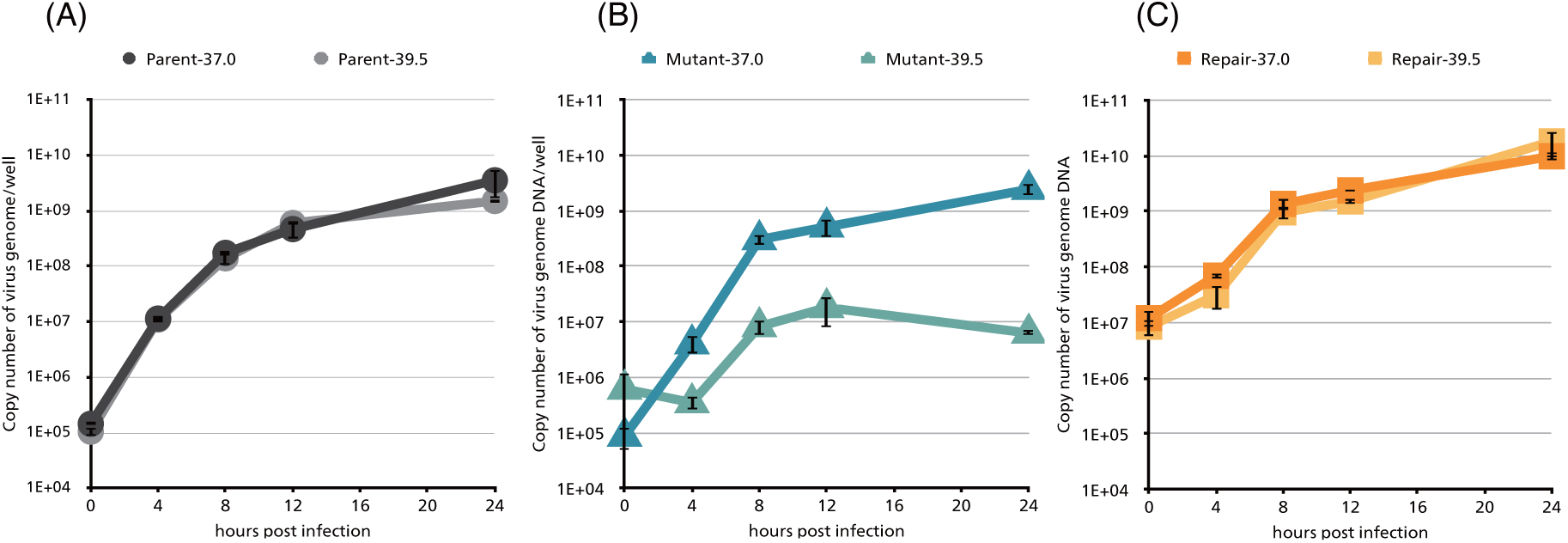
Viral genomic DNA replication kinetics at 37.0℃ and 39.5℃. FHK Tcl3.1 cells were inoculated with virus and incubated at the indicated temperatures. Infected cells were collected and DNA was extracted at 0, 4, 8, 12 and 24 hours post infection. Virus genomic DNA copy numbers were assayed by realtime PCR targeting ORF30. (A) parent Ab4p virus, (B) mutant Ab4p ORF30 N752 virus, (C) repair Ab4p ORF30 D752 virus. The amount of DNA synthesis of the parent and repair viruses were identical at 37.0°C and 39.5℃, while that of the mutant virus at 39.5°C had decreased from 37.0°C.

### Replication capacity of EHV-1 field isolates at 39.5°C

The replication capacities of 49 field isolates were investigated at 39.5°C (Table 2). Virus replication capacity was based on whether a cytopathic effect (CPE) was observed after 72 hours of incubation at 39.5°C. Four EHM-derived isolates (01c1, 89c105, 7759, and 7855) and three non-EHM-derived strains (8343, 9204, and 13757) grew at 39.5°C, while the other 42 non-EHM-derived strains did not.

**Table 2.**
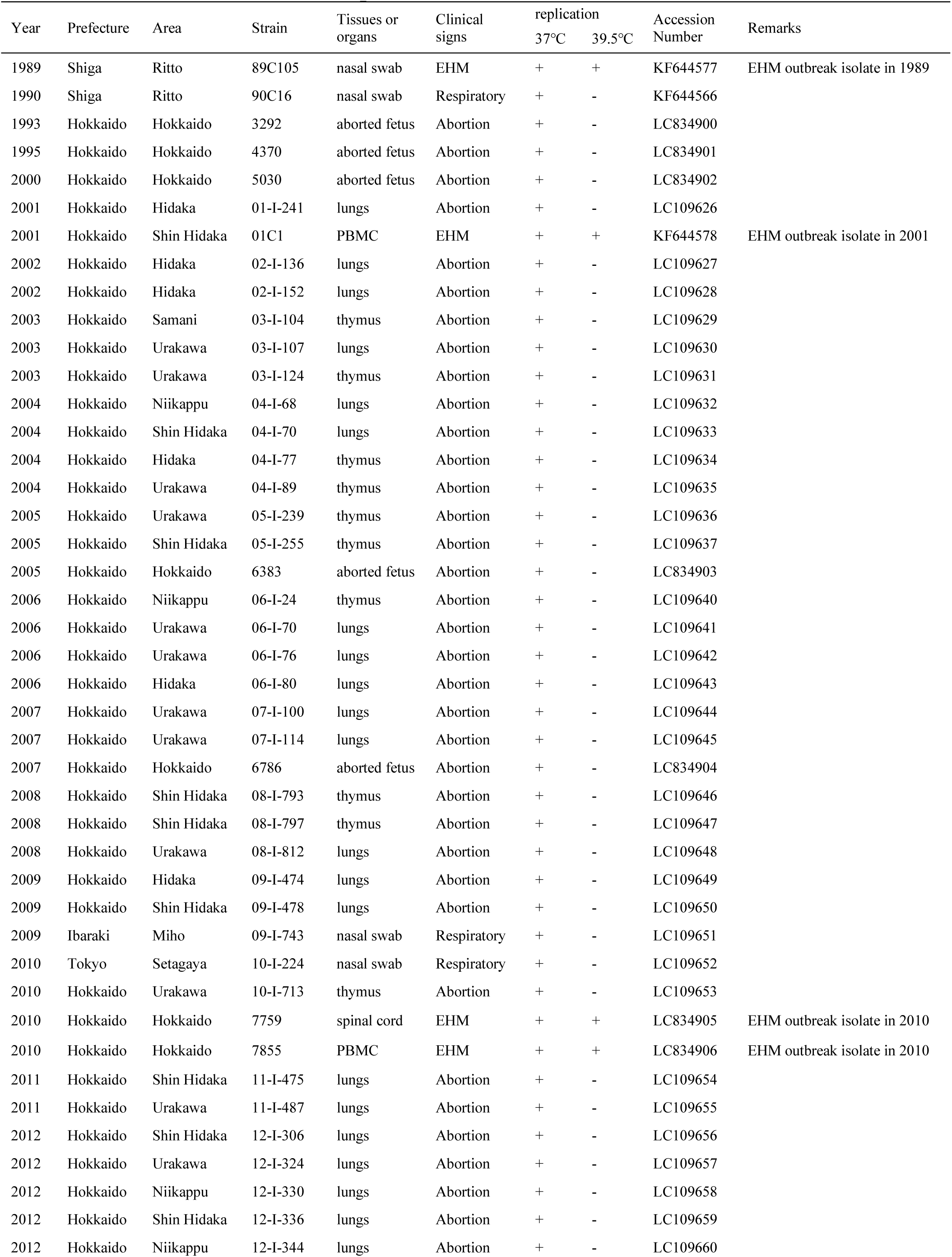
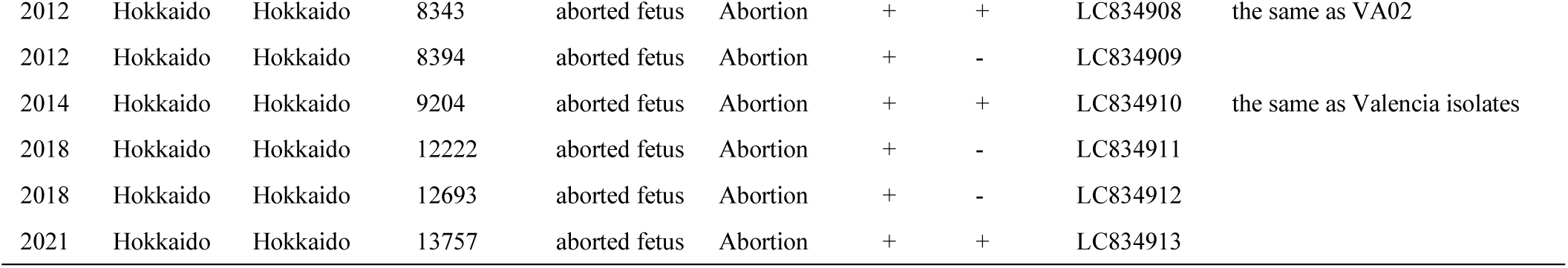
EHV-1 field isolates in Japan.

One of the 39.5°C-grown isolates (01c1) had a D at position 752 in UL30 and six of them (89c105, 7855, 7759, 8343, 9204, and 13757) had an N. Notably, the six N752 isolates had additional SNPs causing other substitutions (A742 of 89c105 and 7855; R250 of 7759; G760 of 8343; I219 of 9204; S151, H625, K683 and T1107 of 13757) (Table 3). The SNP in the 9204 strain (I291) was the same the one in the Valencia strain. Among the 42 strains that did not grow at 39.5°C, two N752 strains had additional SNPs (K683 and N769 of 06-I-70; V946 of 08-I-812). The other 40 strains included 39 strains with N752 and one strain (06-I-80) with D752. 06-I-80 was the only D752 strain that did not grow at 39.5°C.

**Table 3.** Replication at 39.5°C and UL30 amino acid variations of Ab4p, field isolates in Japan and chimera BAC viruses.

| Strains | Diseases | 39.5°C | Amino acids of UL30 |  |  |  |  |  |  |  |  |  |  |
| --- | --- | --- | --- | --- | --- | --- | --- | --- | --- | --- | --- | --- | --- |
|  |  |  | 151 | 250 | 291 | 625 | 683 | 742 | 752 | 760 | 769 | 946 | 1107 |
| Ab4p* | EHM | + | R | H | S | R | E | V | D | D | H | I | A |
| 01c1 | EHM | + | . | . | . | . | . | . | . | . | . | . | . |
| 89c105* | EHM | + | . | . | . | . | . | A | N | . | . | . | . |
| 7855* | EHM | + | . | . | . | . | . | A | N | . | . | . | . |
| 7759* | EHM | + | . | R | . | . | . | . | N | . | . | . | . |
| 8343 | abortion | + | . | . | . | . | . | . | N | G | . | . | . |
| 13757 | abortion | + | S | . | . | H | K | . | N | . | . | . | T |
| 9204* | abortion | + | . | . | I | . | . | . | N | . | . | . | . |
| Valencia** | EHM | + | . | . | I | . | . | . | N | . | . | . | . |
| H752** | EHM | + | . | . | . | . | . | . | H | . | . | . | . |
| 06-I-80 | abortion | - | . | . | . | . | . | . | . | . | . | . | . |
| 06-I-70* | Respiratory disease | - | . | . | . | . | K | . | N | . | N | . | . |
| 08-I-812* | Respiratory disease | - | . | . | . | . | . | . | N | . | . | V | . |

\* Replication at 39.5°C was assayed using the original stock and chimera viruses.
\*\* Replication at 39.5°C was assayed using the chimera viruses.
• Same amino acid as that of Ab4p

### UL30 SNPs other than UL30 D752 and replication capacity at elevated temperatures

We first observed the effects of D752 and N752 on the growth of Ab4p bacterial artificial chromosome (BAC) viruses. The BAC viruses encode a green fluorescent protein (mEGFP) (Appendix Fig. A1), which acts as a marker of viral growth in infected FHK Tcl3.1 cells. At 39.5°C, the parent D752 BAC virus and the D752 repaired BAC virus grew well while growth of the mutant N752 BAC virus was observed in only a few cells (Fig. 4).

**Fig. 4.**
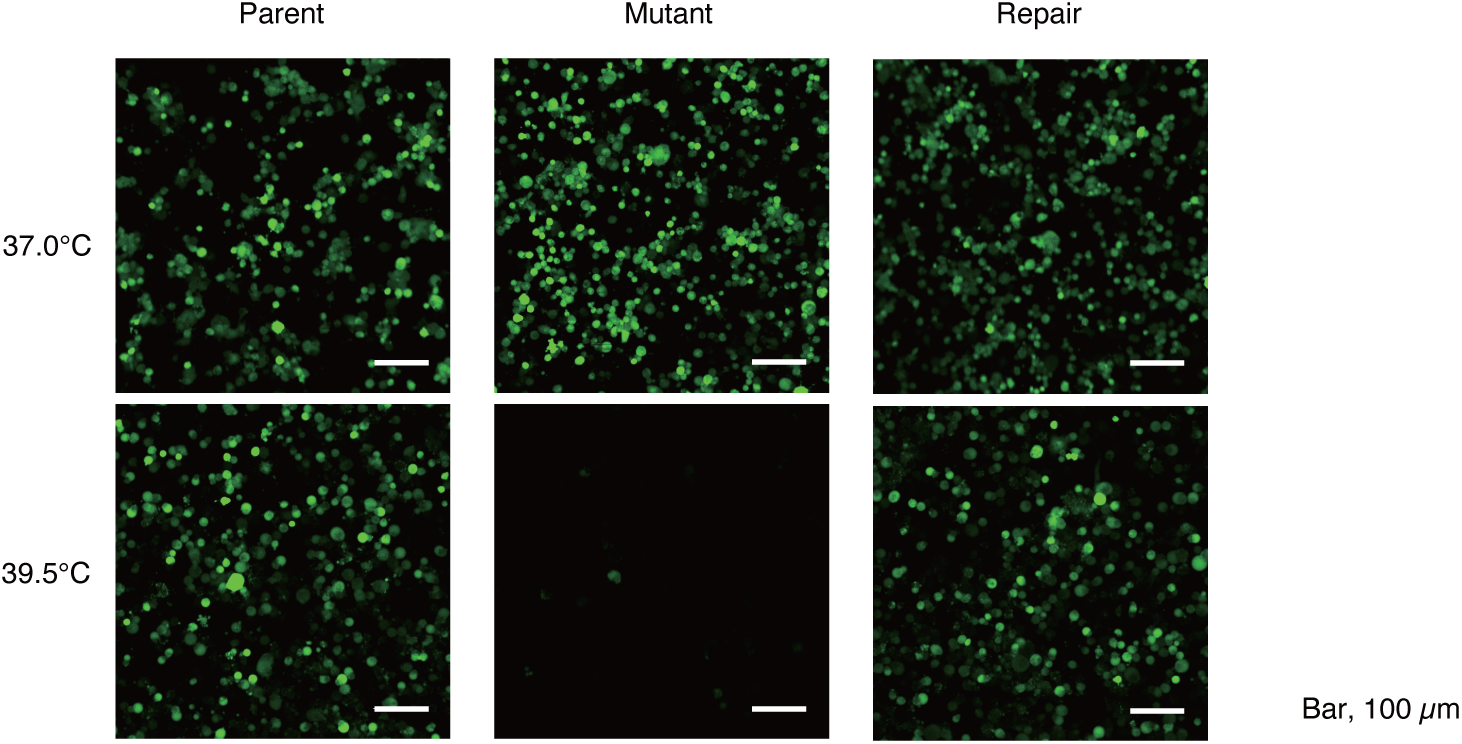
Growth of Ab4p BAC parent, mutant and repaired viruses growth at 37.0℃ and 39.5℃. FHK Tcl3.1 cells were inoculated with BAC viruses and incubated at temperatures indicated in the figure. Fluorescence images were taken at 48 hours post inoculation. BAC vector sequence inserted in the EHV-1 genome includes the mGFP gene which was expressed in infected cells (appendix Fig. A1). Ab4p parent and repair BAC viruses grew at 39.5℃. Mutant (Ab4p ORF30 N752 BAC) did not grow at 39.5℃.

We next investigated the effects of seven other UL30 SNPs (R250, I291, K683, A742, H752, N769, V946) (Fig.5, Table 3) on the growth at 39.5°C. To do this, the SNPs were introduced into individual BAC chimera viruses consisting of an Ab4p background and the ORF30 gene of each virus. Of the seven UL30 SNPs found in the field isolates, four (R250, I291, A742 and H752) enabled growth at elevated temperatures and the other three (K683, N769 and V946) did not (Fig. 5, Table 3).

**Fig. 5.**
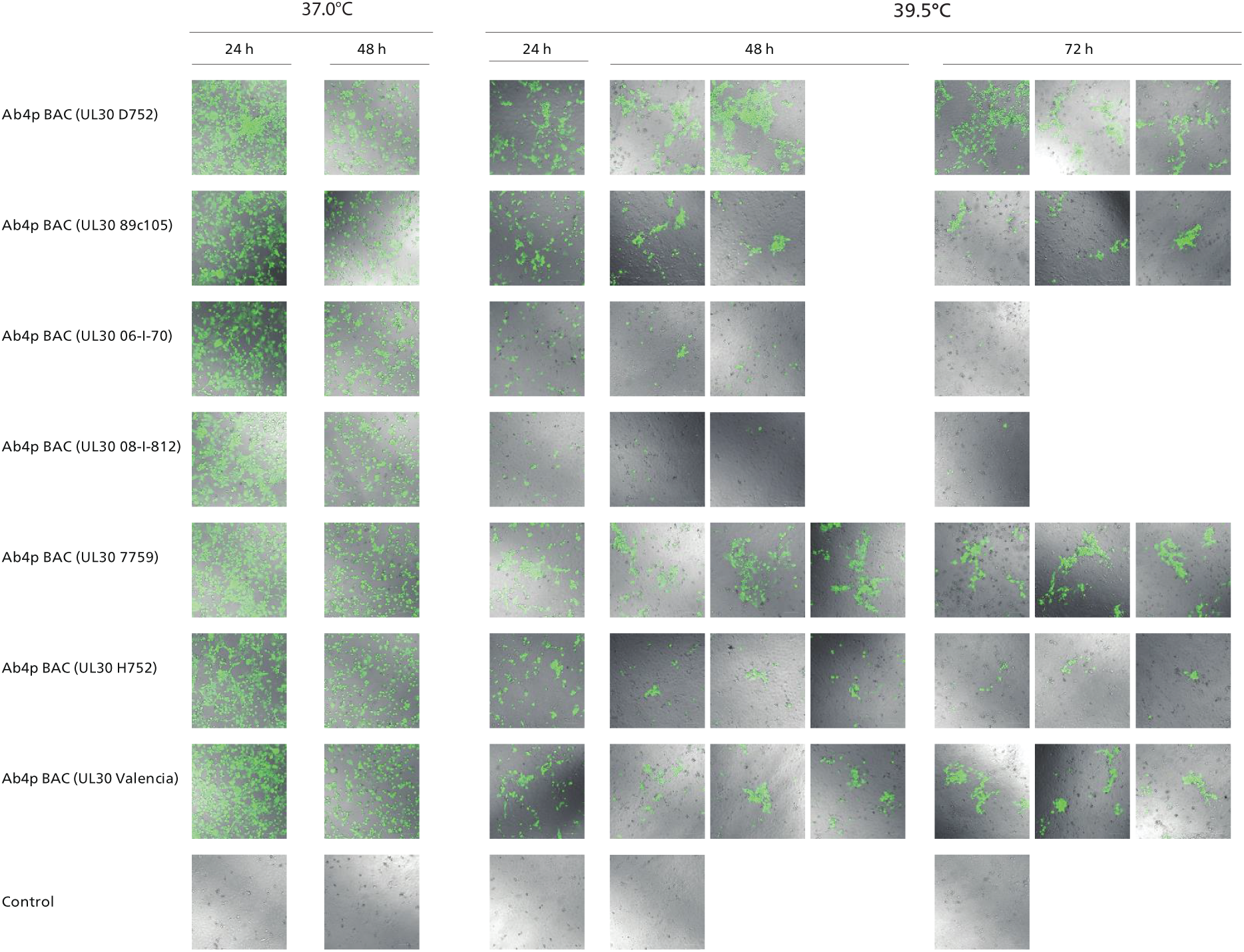
Proliferation of chimera BAC viruses at elevated temperatures. FHK Tcl3.1 cells were transfected with chimera BAC DNAs and cultivated for 72 hours at 37.0°C and 39.5°C. Supernatant of each virus culture was collected and stored at -80°C. FHK Tcl3.1 cells were inoculated with each recovered virus and cultivated for 72 hours at 37 and 39.5°C. Fluorescence images merged with DIC images were taken using a laser confocal microscope at 24, 48 and 72 hours post inoculation.

### SNP mapping in a 3-D model of EHV-1 DNA polymerase UL30

Tertiary UL30 structure models of nine of the variants (Ab4p, 06-I-70, 89c105, 7759, 8343, 13757, H752, V592, and Valencia) were predicted using AlphaFold3. The predicted structures were similar to each other and to the X-ray diffraction-determined structure of the herpes simplex virus -1 (HSV-1) DNA polymerase POL (24), which has 67% similarity to UL30 at the amino acid level. Despite the difference in amino acid sequence, the structures of the functional domains of the UL30s were highly similar to those of the POL domains. The structural domains were named according to Liu et al. (24) (Fig. 6, Appendix Fig. A2). Each of the SNPs in UL30 investigated in the present study was plotted into the 3-D model of EHV-1 Ab4p UL30 (Fig. 6). The SNPs of EHV- 1 UL30 associated with replication capacity at elevated temperatures seemed to centralize in the region adjoining the NH2-terminal and Palm domains of UL30 (the turquoise and blue domains, respectively in Fig. 6). Other SNPs which were not associated with replication capacity at elevated temperatures were scattered in other domains of UL30. SNPs of UL30 associated with replication capacity at elevated temperatures affected DNA polymerase activity but did not appear to affect the 3-D structure of EHV-1 UL30 at elevated temperatures.

**Fig. 6.**
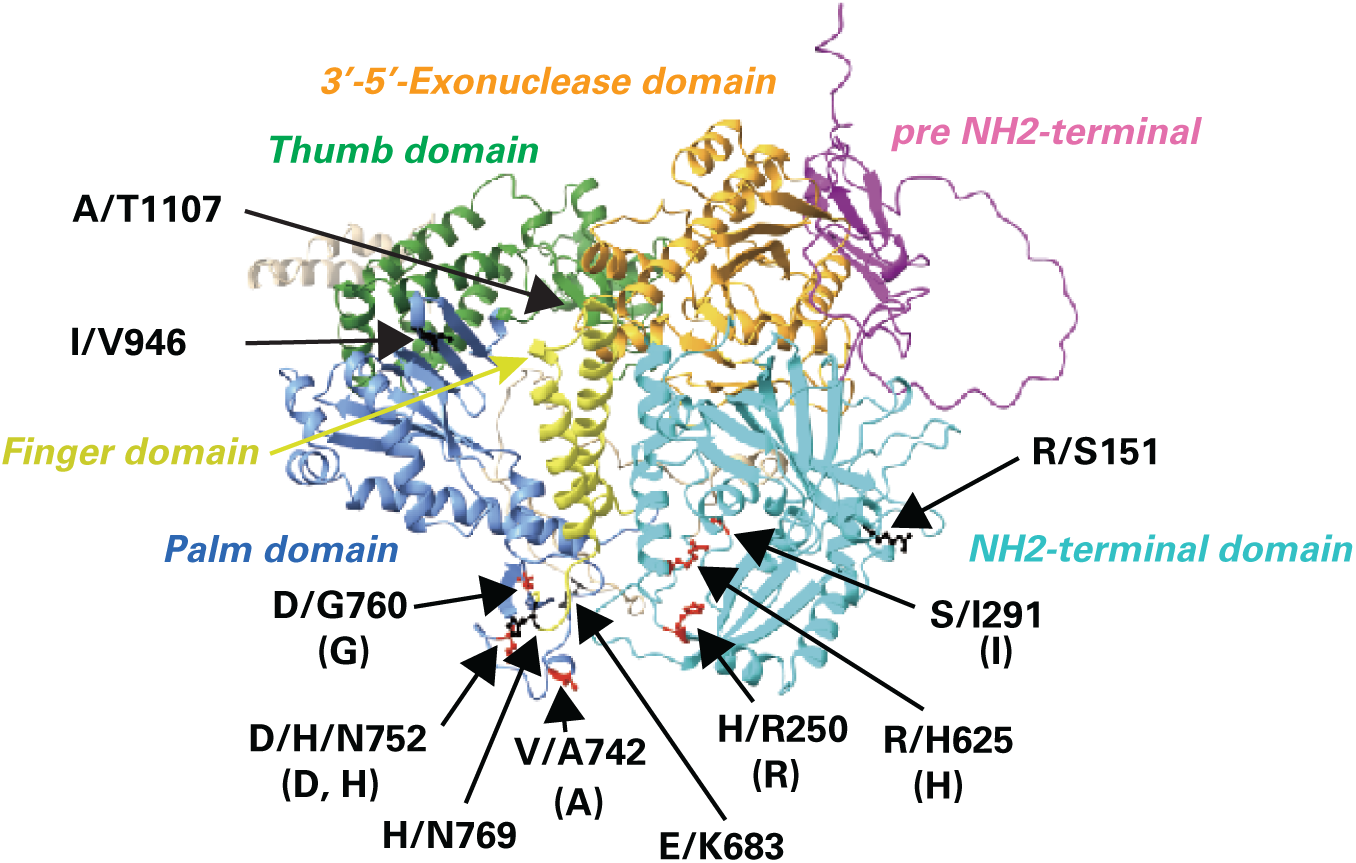
SNPs mapping on 3-D modelling of EHV-1 UL30. Using AlphaFold3, 3-D modelling of EHV-1 Ab4p UL30 D752 was predicted. Functional domains of UL30 are colorized according the 3-D structure of HSV-1 UL30 (24). Amino acids of virus capable of replicating at elevated temperatures are shown in round brackets. Sites of each SNP were mapped on 3-D modelling of EHV-1 Ab4p UL30. EHM associated SNPs (red) are centralized next each other in NH2-terminal (turquoise) (S/I291, H/R250, R/H625) and Palm (blue) (V/A742, D/H/N752, D/G760) domains. SNPs (black) not associated with replication capacity at elevated temperatures are outside of UL30 such as N-terminal domain (R/S151), joining loop between N-terminal and Palm domains (E/K683), Finger domain (H/N769), Palm domain (I/V946), and Thum domain (A/T1107).

## Discussion

D752 was previously identified as a neuropathogenic marker of EHM (14), and in the present study, it was revealed to be associated with replication capacity at elevated temperatures. Furthermore, EHM-derived isolates possessing N752, which is a non-neuropathogenic marker, and other SNPs (e.g., A742 of 89c105 and 7855, and R250 of 7759), showed replication capacity at elevated temperatures. Chimera BAC viruses expressing UL30 SNPs of EHM isolates from other countries were also able to replicate at elevated temperatures. We speculate that EHV-1s that cause EHM can replicate at elevated temperatures whether or not they have D752 in UL30. EHM onset is reported to follow fever in infected horses (8–10). Replication of most of EHV-1strains appear to be repressed or blocked at elevated temperatures in the primary febrile phase of infected horses. However, some strains can replicate under febrile conditions, leading to high virus loads and viremia. Thus, EHV-1s that can replicate at elevated temperatures might be the cause of EHM outbreaks.

Plating efficiency of Ab4p parent and repair viruses did not vary in the range of elevated temperatures examined, although plaque sizes of these viruses decreased at 40.0°C. These results indicated that elevated temperatures retarded the post-entry replication stage of EHV-1-infected cells. The amounts of DNA synthesis of the parent and repair viruses were identical at 37.0°C and 39.5℃, while that of the mutant virus at 39.5℃ was lower than it was at 37.0℃. This indicates that N752 SNP of UL30 decreased DNA polymerase activity at 39.5°C.

SNPs associated with replication capacity at elevated temperatures were centralized in the NH2-terminal and Palm domains (Fig. 6). On the other hand, the amino acid substitutions in UL30 of viruses that did not grow at 39.5°C (K683 and N769 of 06- I-70 and V946 of 08-I-812) were scattered through other parts of the protein. Liu et al. (24) reported that some surface peptides located in the NH2-terminal and Palm domains had very large temperature factors, which is an indication that their structures are very sensitive to temperature. The SNPs in NH2-terminal and Palm regions may be involved in the structural stability needed to function at elevated temperatures. Further studies are needed to understand how viral DNA replication in the mutant Ab4p N752 virus infected cells is suppressed at elevated temperatures.

The results with the 49 Japanese isolates (Fig. 5, Table 3) suggest that the replication capacities at elevated temperatures are largely determined by UL30. The exception (06-I-80) had D752, which suggests that factors other than D752 might be suppressing 06-I-80’s replication at elevated temperatures.

Because all EHM-derived EHV-1s examined in the present study replicated at elevated temperatures, the onset of EHM appears to be related to an ability to replicate at elevated temperatures. Although several Japanese isolates grew at 39.5°C (Table 2), these viruses were all abortion-derived strains and did not cause EHM. Isolate 8343 was isolated from an aborted fetus in 2012, and its UL30 amino acid sequence was the same as that of EHM-derived strain VA02 isolated in Virginia, USA, in 2002. Isolate 9204 was isolated from an aborted fetus in 2014, and its UL30 amino acid sequence was the same as that of EHM-derived strains of Valencia outbreak in 2021. The ORF30 SNPs associated with replication capacity at elevated temperatures, including the one at position 2279 (an A-to-G nucleotide substitution) causing a D760G amino acid substitution in the Japanese isolates, have been found in other countries. Nugent *et al.* (14) reported D760G substitutions in an EHM strain (BE95_1_2, US99_3_2, US02_1_2) and in an abortion-derived strain (GB04_8_1). Host factors might also be involved in EHM but four genome-wide association studies failed to find any (13, 25, 26, 27). We cannot rule out the possibility that viral and host factors other than UL30 have a role in the pathogenesis of EHM.

The present results reveal that the previously found neuropathogenic marker, D752, and other SNPs found in ORF30, viral DNA polymerase gene, are related to replication capacity at elevated temperatures. This finding may be useful in the search for EHM countermeasures.

## Materials and Methods

### Cells

Fetal horse kidney-derived FHK Tcl3.1 cells were propagated in Dulbecco’s modified Eagle’s medium (DMEM) (FUJIFILM Wako Pure Chemical Corporation, Osaka, Japan) containing 10% fetal bovine serum (FBS) (28). The cells were maintained in a humidified 5% CO_2_ atmosphere at 37°C.

### Viruses

EHV-1 Ab4p strain was kindly provided by Dr. A. J. Davison, Glasgow University, Scotland (29, 30). A bacterial artificial chromosome (BAC) vector, pZC320mGFP, was inserted into the virus genome of Ab4p to construct pAb4p BAC (a parental BAC plasmid) in our laboratory as described previously (Appendix Fig. A1) (23). pZC320mGFP was constructed based on a mini-F plasmid pZC320 obtained from NBRP E.coli Strain (National Institute of Genetics, Mishima, Japan) to contain a modified green fluorescent protein (mGFP) gene as one of the selection markers (23, 31). Using pAb4p BAC, a single nucleotide substitution at 2254 site of UL30 was introduced to construct a mutant BAC, pAb4p UL30 N752 BAC, in which the 752nd amino acid codon at UL30 was replaced from aspartic acid codon (GAC, D752) to asparagine codon (AAC, N752), and a repair BAC, pAb4p UL30 Repair BAC, in which 752nd codon was back to the original aspartic acid codon (GAC, D752). Viruses were recovered as recombinant viruses without BAC including Ab4p UL30 (a parent virus), Ab4p UL30 N752 (a mutant virus), and Ab4p UL30 Repair (a repair virus) and recombinant viruses with BAC including Ab4p UL30 BAC (a parent BAC virus), Ab4p UL30 N752 BAC (a mutant BAC virus) and Ab4p UL30 Repair BAC (a repair BAC virus) as described previously (23).

Forty-nine EHV-1 field isolates in Japan including four EHM isolates (89c105, 01c1, 7759 and 7855) were shown in Table 2. Viruses used in the present study were propagated in FHK Tcl3.1 cells in a 5% CO_2_ cell culture incubator at 37°C. Full genome sequences (KF644566, KF644577, KF644578, LC109626 to LC109660, LC834905, LC834906) and ORF30 sequences (LC834900 to LC84904, LC834907 to LC834913) of the field isolates in Japan have been registered into DDBJ (Table 2) (a manuscript in preparation).

### Evaluation of replication capacity of viruses at elevated temperatures

Parent, mutant, and repair viruses without BAC vector sequence were subjected to plaque formation assay at elevated temperatures. Using two cell culture incubators, of which the interior was maintained with a humidified 5% CO_2_ atmosphere, one was set at 37.0°C (MCO-170A/CUVD-PJ, Panasonic, Tokyo, Japan), and the other was set at 38.5°C, 39.0°C, 39.5°C, 40.0°C, or 40.5°C (CPO2-2301, KK-Hirasawa, Tokyo, Japan). The temperature fluctuation was ±0.1°C. In brief, confluent FHK Tcl3.1 cells were prepared in duplicate 24-well plates at 37.0°C. Serial dilutions of the stock virus (10^7^pfu/ml) were made with DMEM. FHK Tcl3.1 cells were inoculated with 10^-4^, 10^-5^, 10^-6^ dilutions of virus and incubated at 37.0°C and 38.5°C. For the mutant virus at 39.0°C and higher temperatures, 10^-1^, 10^-2^, 10^-3^ dilutions were used. One hundred µL/well of diluted virus samples were inoculated into FHK Tcl3.1 cells. Each dilution of the virus was inoculated into four wells. After one hour adsorption at 37.0°C, inoculated solution was removed, and cells were overlaid with 0.5 mL/well of DMEM containing 1% methylcellulose and 1% FBS. One plate was incubated at 37.0°C and the other was at an elevated temperature. After 3 days, 0.5 mL/well of 0.15% crystal violet in 20% ethanol was directly added for staining cells for 3 days, washed with tap water and dried. Plate images were taken using a digital camera. Using the image files, plaque areas were measured with ImageJ 1.54M (http://imagej.org). Plaque areas were statistically analyzed using Wilcoxon rank sum test in RStudio (Version 2024.12.1_563, Posit Software, PBC). Graphs were drawn using ggplot program in RStudio and labels were edited using Adobe Illustrator 2025 (Adobe, SanJose, USA).

### Viral growth kinetics

Monolayers of FHK-Tcl3.1 cells in 4-well plates were inoculated with viruses at a multiplicity of infection (MOI) = 0.1. Virus stock solution, which was added DNase I (1 µL for 1.5 mL virus stock solution) and incubated for 15 min at 37°C, was inoculated to FHK Tcl3.1 cells. After one hour adsorption, the inoculum was removed and added 0.5 mL/well of DMEM containing 1% FBS was added. The inoculated cells were incubated at 37.0°C or 39.5°C. Supernatants and cells were collected at 0, 4, 8, 12 and 24 hours post inoculation. The culture medium was collected, centrifuged, and the supernatant was stored at -80°C. Infected cells remained in wells were washed three times with PBS, once with 0.25% trypsin-0.02% EDTA (Sigma-Aldrich Japan, Tokyo, Japan), incubated for 5 min and suspended in 0.5 mL PBS. Centrifuged at 12,000 rpm for 1 min at 4°C, the supernatant was removed, and the cell pellet was resuspended in 0.5 mL PBS, and stored at -80°C.

### Virus titration

Virus supernatant samples were subjected to plaque formation assay. In brief, confluent FHK Tcl3.1 cells were prepared in a 24-well plate. Serially diluted virus samples were inoculated into FHK Tcl3.1 cells. Four wells per dilution were inoculated. After one hour adsorption, inoculated solution was removed, and cells were overlaid with 0.5 mL/well of DMEM containing 1% methylcellulose and 1% FBS. After 3 days, the cells were stained with adding an equal volume of 0.15% crystal violet in 20% ethanol for 3 days, washed with tap water and dried.

### DNA extraction

DNA was extracted from infected cells using SepaGene (Sekisui Medical Co., Ltd. Tokyo, Japan) according to the manufacturer’s protocol. DNA extracted was dissolved in TE (10 mM Tris-HCl, pH 8.0, 1 mM EDTA) and stored at -20°C until use. DNA concentration was measured with a Qubit 2.0 Fluorometer (Thermo Fisher Scientific K.K., Tokyo, Japan) using a Qubit dsDNA BR Assay kit.

### Realtime PCR

DNA copy numbers were titrated with Thermal Cycler Dice Real Time System (TaKaRa Bio, BIO, Kusatsu, Japan) using TB Green Premix Ex Taq II (Tli RNaseH Plus) (TaKaRa Bio, Bio, Kusatsu, Japan). Primers for the realtime PCR targeting ORF30 gene were UL30F, 5’-gtcaggcccacaaacttgat-3’, and UL30R, 5’-actcggtttacggattcacg-3’. PCR program was 95°C for 30 sec of initial denaturation, 40 cycles of two steps, 95°C for 5 sec and 60°C for 30 sec, followed by dissociation step of 95°C for 15 sec, 60°C for 30 sec and 95°C for 15 sec. Purified pAb4p BAC DNA was used as a reference DNA sample for quantitation.

### Construction of chimera BAC viruses

We constructed chimera BAC viruses containing nucleotide mutations in ORF30 gene, which mutations were found in ORF30 gene as SNPs (T2225C/V742A and G2254A/D752N; G2047A/E683K, G2254A/D752N and C2305A/H769N; and C748G/H250R and G2254A/D752N, G2254A/D752N and A2836G/I946V) of field isolates in Japan, SNPs (G872T/S291I and G2254A/D752N) of one field isolate in Japan and Valencia strains and a new SNP (G2254C/D752H) of isolates in US and Europe, based on pAb4p BAC and pAb4p ΔORF30 rpsLneo BAC as parent BACs using the BAC technology established in our previous study (23, 31) and another BAC plasmid, pAb4p ORF30 Δaa55-494_rpsLneo BAC, which was constructed in the present study as follows. A rpsLneo cassette with arms was amplified using a rpsLneo DNA as a template with primers Δ30(55-494)rpsL-neo F: 5’- tcaccttgctcttcttctgaaaatggttcgtggcgatgtcccacaccttGGCCTGGTGATGATGGCG-3’ and Δ30(55-494)rpsL-neo R: 5’- gcccaaggcccccccaacactcgtactgcacagaggtgggtagctttaagTCAGAAGAACTCGTCAAGAA GGCG-3’ and PrimeSTAR MAX DNA polymerase (TaKaRa Bio, Bio, Kusatsu, Japan). In the primer sequences, the arm sequences are shown in lower case and homologous sequence to rpsLneo are shown in upper case. The rpsLneo cassette with arms was electroporated into *Eshcerichia coli* DH10beta containing pAb4p BAC and pRED/ET plasmids. The rpsLneo cassette was replaced with a part of ORF30 gene using RED/ET gene replacement system, Counter Selection BAC Modification Kit (Gene Cambridge GmbH, Heidelberg, Germany). These BACs contained pZC320mEGFP as the BAC vector sequence. pZC320mEGFP encodes the fluorescent protein mEGFP gene as a marker (31).

Site directed mutagenesis for G872T/S291I and G2254C/D752H in ORF30 was examined by inverse PCR with primers containing desired mutations for each site. In brief, a fragment of ORF30 containing A2254 mutation (23) was amplified using a pair of primers, infusion ORF30 pU19F 5’-acgaggccctttcgtcggccacatcctcgtcgtaca-3’ and infusion ORF30 pUC19R 5’-caatacgcaaaccgcacgggtacggcaagttcaac-3’, Ab4p UL30 N752 DNA as a template and PrimeSTAR Max DNA polymerase (TaKaRa Bio, Bio,, Kusatsu, Japan) with an amplification program of 30 cycles of 98°C for 10 sec, 60°C for 5 sec and 72°C for 8 sec. The PCR product was purified using NucleoSpin (TaKaRa Bio,, Kusatsu, Japan). DNA concentration was measured using Qubit 2. The UL30 fragment was inserted into pUC19 using In-Fusion system according to the manufacturer’s protocol (TaKaRa Bio,, Kusatsu, Japan). In-Fusion solution was transformed into *E. coli* DH5alpha Competent Cells (TaKaRa Bio,, Kusatsu, Japan) and plated on agar plates containing 50 µg/mL ampicilin. The plasmid cloned was designated pUC-UL30. A nucleotide mutation was introduced into pUC-UL30 using PrimeSTAR mutagenesis kit (TaKaRa Bio, Bio, Kusatsu, Japan) with primers of ORF30 G872T F 5’- gaggggaTcgtggacgtgaccacgcg-3’ and ORF30 G872T R 5’-gtccacgAtcccctcgaattttgtaatc- 3’ for G872T (S291I) mutation or primers of ORF30_G2254C_F 5’- gagtagtggacggatggttgaagccc-3’ and ORF30_G2254C_R 5’-tccgtccactactcgacgttcgagg-3’ for G2254C (D752H) mutation. Purified plasmid DNA was sequenced whether the desired mutation had been introduced.

A PCR fragment of ORF30 was amplified using each of DNA extracted from field isolates (89c105, 06-I-70, 08-I-812) and a mutagenized plasmid DNA for G2254C (D752H) as a template DNA with primers ΔrpsL ORF30 F: 5’-ggccacatcctcgtcgta-3’ and ΔrpsL ORF30 R: 5’- tttaaggtgtgggacatcg-3’ and PrimeSTAR Max DNA polymerase (TaKaRa Bio,, Kusatsu, Japan). A PCR fragment of another part of ORF30, a 62 to 1575 region corresponding to amino acid sequence 55 to 494 of UL30, was amplified using EHV-1 7759 DNA or a mutagenized plasmid DNA for G872T (S291I) as a template DNA with primers ORF30 62-82-F: 5’-gcaagaggccatttttcaggc-3’ and OR30 1575-1555- R: 5’-cgtcgccacagaatacatgtc-3’ and PrimeSTAR max DNA polymerase (TaKaRa Bio,, Kusatsu, Japan). PCR products were purified by NucleoSpin (TaKaRa Bio,, Kusatsu, Japan). DNA concentration was measured with a Qubit 2.0 Fluorometer (Thermo Fisher Scientific K.K., Tokyo, Japan) using a Qubit dsDNA BR Assay kit.

Each of the PCR fragments was introduced into *E. coli* DH10beta containing pAb4p ΔORF30 rpsLneo BAC or pAb4p ORF30Δaa55-494_rpsLneo BAC, and pRED/ET plasmid. The rpsLneo cassette in the BAC plasmid was replaced with a ORF30 fragment introduced desired mutation using Red/ET gene replacement system, Counter Selection BAC Modification Kit (Gene Cambridge GmbH, Heidelberg, Germany). Desired mutations introduced in ORF30 of each BAC plasmid DNA were confirmed by Sanger sequencing. The chimera BAC DNA was digested with EcoRI, HindIII or SalI. The digested DNAs were analyzed by 0.5% agarose gel electrophoresis in 40 mM Tris-acetate, 1mM EDTA, pH8.3. The gel slabs were stained with 0.5 µg/mL Ethidium Bromide. The fluorescence image was taken for analysis. No large deletion/insertion was confirmed in the chimera BAC DNA (Appencies Fig. A3).

Each recombinant chimera BAC virus was recovered as previously described (31). Briefly, 200 ng BAC DNA, which was purified using NucleoBond BAC100 (TaKaRa Bio, Bio, Kusatsu, Japan), was transfected into FHK-Tcl3.1 cells seeded in 24- well-plates by using Lipofectamine 3000 (ThermoFisher Scientific K.K., Tokyo, Japan). Transfected cells were then cultured at 37°C for 3 to 5 days. After confirming that the cells expressed green fluorescence and showed a cytopathic effect (CPE), the supernatant was harvested and used to inoculate new cells. The recovered virus was subjected to three rounds of plaque purification. Replication of recombinant chimera BAC viruses was measured at 37.0 and 39.5°C as described above.

### Laser scanning microscope observation

Cells were observed with a confocal laser scanning microscope (Zeiss LSM700; Carl Zeiss, Tokyo, Japan). Zeiss Zen 2009 software was used for the acquisition of images.

### Protein Structure Predictions

The EHV-1 UL30 protein 3-D structure model was predicted using AlphaFold 3 (https://alphafoldserver.com) (32) using the default parameters. The predicted models were visualized with the open-software, UCSF ChimeraX 1.9 (https://www.cgl.ucsf.edu/chimerax/) (33). The predicted protein structures were aligned and superimposed using MatchMaker tool of Chimera-X. Structural domains and secondary structure assignments were assigned onto Ab4p UL30 amino acid sequence according to Liu et al. (24) based on a pair-wise alignment of amino acid sequences of HSV-1 KOS UL30 (accession: JQ673480) and EHV-1 Ab4p UL30 (accession: AY665713.1) using AliView 1.30 (https://github.com/AliView/AliView) (34) (Appendix Fig. A2).

## Acknowledgments

We acknowledge generous support from Prof. T. Asai, Gifu University, and Prof. T. Koshizuka, Gifu Pharmaceutical University. We acknowledge NBRP-E.coli Strain at National Institute of Genetics, Mishima, Japan, for providing us the plasmid pZC320. Funding: Japan Society for the Promotion of Science JSPS KAKENHI Grant Number JP18H02343 and 22K06024 (HF).

## Author contributions

Conceptualization: HF, NF

Methodology: HF, RK, NF, FN, KT

Investigation: HF, RK, NF, FN, KT

Visualization: HF, RK, NF, FN, KT

Funding acquisition: HF

Project administration: HF

Supervision: HF, RK, NF, KT

Writing – original draft: HF, NF

Writing – review & editing: HF, RK, NF, FN, KT

## Data and materials availability

All data are available in the main text or the supplementary materials.

## Appendices

